# Hsp22 is the key sensor and balancer in mitochondrial dynamic associated metabolic reprogramming

**DOI:** 10.1101/2020.11.22.393116

**Authors:** Zaiwa Wei, Ying Cui, Xueyi Wen, Wenjing Wang, Jingxin Mo, Rujia Liao, Yanmei Hu, Quan Zhou, Fang Shi, Tianchan Peng, Ning Tian, Yafang Tang, Lili Wei, Liangxian Li, Xiaoli Wei, Qinghua Li

## Abstract

Altered mitochondrial dynamics are commonly seen in tumors and Parkinson’s disease (PD), but the exact mechanism is unclear. We observed in tumor and PD flies, interference of mitochondrial fission or fusion reverses metabolic reprogramming and attenuates pathogenic phenotypes, indicating the rebalanced mitochondrial dynamics can re-establish cellular homeostasis. Surprisingly, we found Hsp22 is highly induced in tumor and PD flies. Hence, Hsp22 overexpression, not only suppressed RAS tumor flies, hepatocellular carcinoma mice, but also rescued PINK1^B9^ PD flies associating with restored mitochondrial homeostasis and reversing metabolic reprogramming. We speculate that mitochondrial dynamics is normally in a constant dynamic equilibrium, as if on a continually oscillating seesaw, tumor and PD are like the two extreme states of the seesaw which can be corrected by the promising target, Hsp22.

**One Sentence Summary:** Mitochondrial dynamics defects including tumor and PD lead to metabolic reprogramming accompanied with Hsp22 induction, vice versa, overexpression of Hsp22 reverses the mitochondrial dynamic dysfunction caused pathogenic metabolism.

## Introduction

When mitochondrial quality control (MQC) is disturbed, mitochondrial morphodynamics would be impaired which is widely thought to trigger various mitochondria defects related diseases associated with onset of tumor and neurodegenerative disorders [1]. It is well recognized the crucial linkage of mitochondrial dynamics with tumor progression, for example hepatocellular carcinoma (HCC), and there already has studies proved enforcing mitochondrial fission inhibited migration, invasion, and metastasis in human breast tumor [2,3]. However, it is still unclear how metabolism alters with mitochondrial dynamics interference in tumor models. What’s more, dysfunction in mitochondrial dynamics has been linked to neuropathies and is increasingly being linked to neurodegenerative diseases, among them, loss of function mutations in PINK1 cause early-onset autosomal recessive PD which has been highlighted in pathogenesis of mitochondrial dysfunction and evokes the activation of metabolic reprogramming [4]. The similar mitochondrial dynamics defects initiation both ended up in metabolic reprogramming in tumor and PD which brought us the hypothesis that there might be a common sensor and balancer that could be a promising therapeutic target for mitochondria associated metabolic diseases.

Small heat shock proteins (sHsps) are among the most widespread molecular chaperones ubiquitously distributed in numerous species, from bacteria to humans [5]. Among them, Hsp22 participates in the regulation of proteolysis of unfolded proteins that would be upregulated under heat or toxic stress [6,7]. Besides, Hsp22 has been proved to be directly or indirectly involved in development of tumor, neurodegenerative diseases and aging [8]. Here, we investigated the molecular functions of Hsp22 in relation with mitochondrial dynamics defects associated metabolic reprogramming which may shed a light on the uncovering mystery between tumor and PD and the potential therapeutic target on mitochondrial dynamics defects’ diseases.

## Results

### 1 Mitochondrial dynamic balance disturbed models including tumor and PD present metabolic reprogramming accompanied by Hsp22 induction (Fig1 and FigS1)

Dysregulated metabolism is a common feature of the metabolic diseases, including obesity, diabetes, and tumor, which can be regulated by mitochondrial dynamics. To investigate whether arbitrarily modulation of mitochondrial dynamics alters metabolic programming, we used *MHC-Gal4* to muscle-specifically drive the genetically modified genes that most influential on mitochondrial morphology including the GTPase dynamin-related protein 1 (Drp1) overexpression (OE), Mitochondrial associated regulatory factor (Marf) RNA interference (RNAi), Marf OE which were confirmed by RT-PCR on the mRNA expression except Drp1 dominant negative (DN). In tumor, the cell over-proliferation model, mitochondria are important mediators of tumorigenesis, as this process requires flexibility to adapt to cellular and environmental alterations in addition to tumor treatments such as chemotherapy. In PD which can be regarded as the cell reduction model is caused by the death or malfunction of dopaminergic neurons leading to muscle movement and coordination impairment were detected with the impaired mitochondrial dynamics and function. More interestingly, the two oppositely developed cell proliferation models, tumor and PD were widely thought to be rarely existing in the same pathogenesis. Moreover, we found that with mitochondrial dynamics modulation, flies’ motor activity phonotypes altered significantly compared with W1118 controls including abnormal wing posture and flying ability (Figure 1A). With the detection of TEM, we found the increased fission capacity of mitochondria in adult indirect flight muscles of Drp1 OE and Marf RNAi MHC-Gal4 driven flies while Drp1 DN and Marf OE flies presented sever fused mitochondrial shapes (Figure 1B) further demonstrates the crucial role of mitochondrial morphology in the determination of the motor function. Ubiquinone Oxidoreductase Core Subunit S3 (*NDUFS3*) is an iron–sulfur subunit which encodes the seventh and last constitutive subunit of the CI core and plays an important role in maintaining membrane potential and ATP generation. We confirmed NDUFS3 expression levels in mitochondrial dynamic modulated flies by immunoblot, consistently we found that corresponds to ETC CI function *NDUFS3* expression decreased in motor activities decreased, tumor and PD flies (Figure 1C, Figure S1B and Figure S1G, Figure S2A, B and C). Mitochondrial dysfunction particularly decreased the function of the electron transport chain (ETC), mitochondrial Complex I (CI) and Complex II (CII) are often regarded as the most likely sites of an ETC impairment. Thus, we conducted oxygen consumption experiments on mitochondrial dynamics altered flies’ thorax tissue as previously described. Interestingly, we observed the dramatically reprogrammed CI and CII functions compared with W1118 controls in mitochondrial dynamics modulated flies while they decreased significantly in tumor and PD fly models (Figure 1D, Figure S1B and S1G). Since mitochondrial respiration and glycolysis are two major energy-yielding pathways, we screened glycolysis signature genes to detect glycolysis metabolism during mitochondrial dynamic alterations. Surprisingly, we observed the dramatically altered Pglym78, PGK, PYK, PFK, GAPDH1 and GAPDH2 mRNA expression and the end product of glycolysis, lactate levels in mitochondrial dynamics arbitrary regulated flies (Figure 1E and F) including tumor and PD flies (Figure S1C, D and S1H, I).

**Figure 1.**
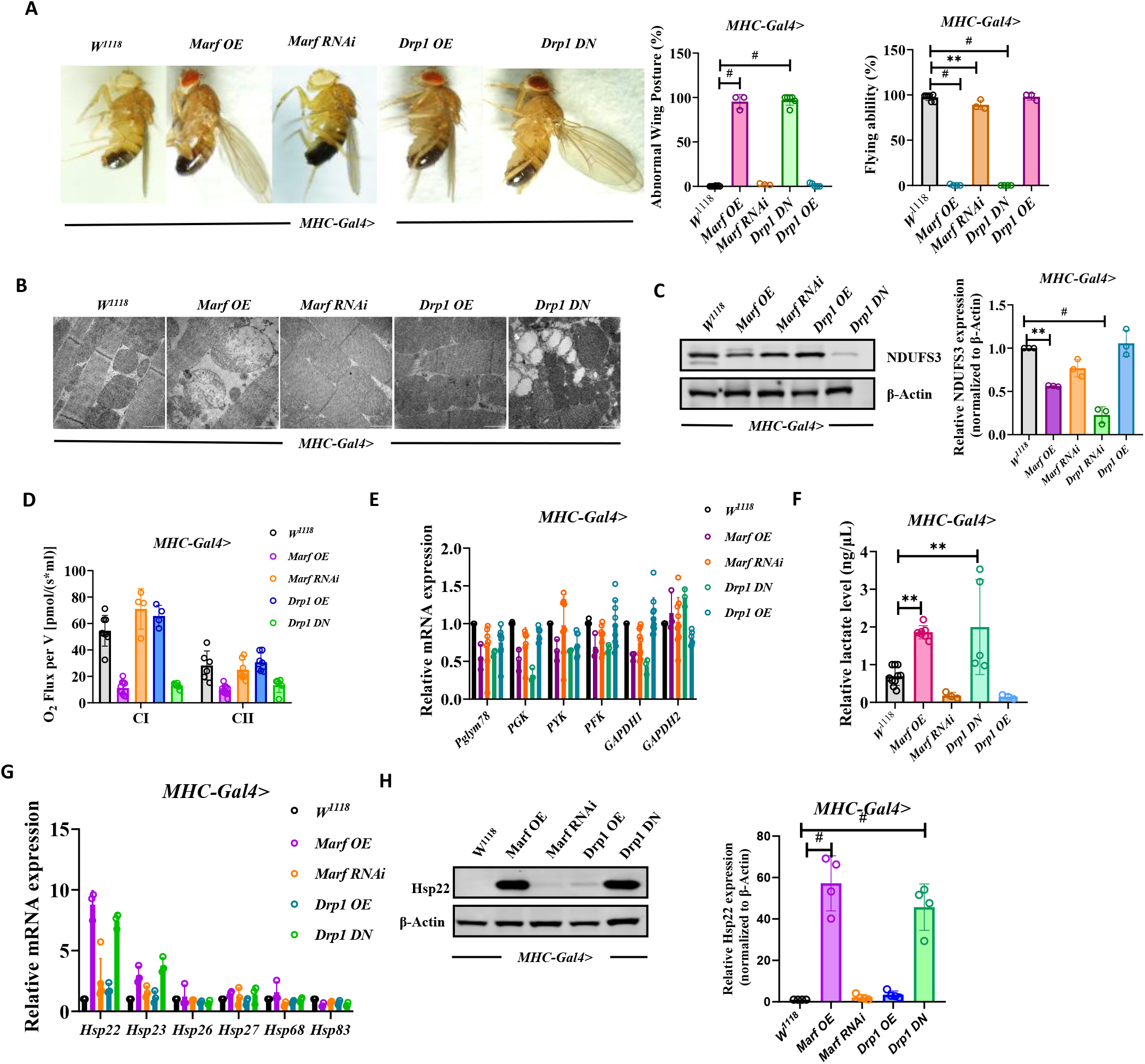
Arbitrarily regulated mitochondrial dynamic models present metabolomic reprogramming accompanied by Hsp22 induction. (**A**) *MHC-Gal4* driven mitochondrial dynamics modulated fly motor actives. (**B**) Mitochondrial morphology changes tested by TEM on arbitrarily regulated mitochondrial dynamics. (**C**) *NDUFS3* protein expression tested by WB separately in each group. (**D**) Quantification of mitochondrial CI and CII function tested in thorax tissue. (**E**) Relative glycolytic marker of *MHC-Gal4* driven mitochondrial dynamics modulated mRNA expression. (**F**) Relative lactate level tested separately in each group. (**G**) Screenings of essential Hsp mRNA expression via RT-PCR. (**H**) *Hsp22* protein expression tested separately in each group. All values are means ± SD of at least three independent experiments. Student’s t test (unpaired); *p< 0.05, **p< 0.01, **p< 0.001.

The common sensor and balancer during mitochondrial dynamic alteration determined metabolic reprogramming that connects inside of mitochondrial metabolism and outside in cell matrix remains a mystery since members of the Hsp family are mainly responsible for proper protein folding and performing many other functions in living organisms such as some of them present in mitochondria and cytosol of eukaryotic cells, as well as on their surface. Here we conducted RT-PCR screening on Hsps to confirm which Hsp is mostly induced in response to exposure to mitochondrial dynamics alteration (Figure 1G). We observed the significantly increased Hsp22 mRNA expression in mitochondria dynamics modulated flies which was also found in tumor and PD flies while there were no obvious expression level changes in other Hsps (Figure 1G, Figure S1E and J). As expected, Hsp22 protein levels increased significantly consistent with mRNA level changes in mitochondrial dynamics modulated flies compared with W1118 controls (Figure 1H).

### 2 Tumor Drosophila model could be rescued by modulation of mitochondrial dynamics with restored Hsp22 expression

To investigate whether mitochondrial dynamics modulation could impact on RasV12; Scrib-/- flies, we directly interfered tumor flies with Drp1 OE, Marf RNAi, Drp1 DN and Marf OE genetically. Surprisingly, we found that no matter fission or fusion trended modulation both suppressed RasV12; Scrib-/- flies’ tumor invasion (Figure 2A) even with arbitrarily modulation of PINK1 and Parkin (Figure S3A) while PINK1 and Parkin are related to autosomal recessive PD and MQC. Since proto-oncogenes Ras, Myc activation collaborate with HIF1-α to confer metabolic reprogram in tumor cells. The suppression effect on the clones of RasV12; Scrib-/- flies was further confirmed by RT-PCR from which we observed the decreased Ras, Myc and Hif1-α mRNA levels (Figure 2B and Figure S3B) which is consistent with the inhibition of the tumor invasion (Figure 2A and Figure S3A). In tumor cells, the conversion of pyruvate into lactate takes place even in the presence of oxygen which is called the ‘Warburg Effect’. Hence, we tested glycolysis signature genes to see how the rescued tumor flies reprogrammed metabolism. Interestingly, we found that PFK, GAPDH1, GAPDH2, PGK, Pglym78 and PYK mRNA in glycolysis pathway decreased significantly in the rescued arbitrarily modulated mitochondrial fission and fusion dynamics even in PINK1 and Parkin interfered RasV12; Scrib-/- tumor flies with the decreased lactate levels (Figure 2F and G; Figure S3G and H). Furthermore, mitochondrial function was examined as it’s the second important energy source during tumor pathogenesis. As expected, we found that in mitochondrial dynamics modulated RasV12; Scrib-/- tumor flies showed the improved ETC CI and CII functions with the increased NDUFS3 expression (Figure 2D and E; Figure S3E, F). Surprisingly, we found that Hsp22 mRNA level and Hsp22 protein expression restored even like the W1118; Scrib-/-control flies which is opposite to the RasV12; Scrib-/- tumor flies whose Hsp22 was induced severely no matter in mitochondrial dynamics regulated or PINK1, Parkin interfered flies (Figure 2H and Figure S3I). On the one hand, we noticed that Hsp22 might be a sensor that is sensitive to metabolism disturbed models, which would be restored to normal expression level when the metabolism is balanced; on the other hand, we were surprised to see in RasV12; Scrib-/- tumor flies both Drp1, and Marf mRNA expression increased dramatically indicating there might exist a highly pathogenically balanced mitochondrial dynamics and any disturbance of this dynamics including PINK1 and Parkin interferences rescued its pathogenic metabolism leading to a reprogrammed relatively healthy status which assumes that the highly pathogenically balanced upregulation of biosynthetic and bioenergetics pathways especially glycolysis and mitochondrial biogenesis was destroyed which brought the cellular metabolism to a relatively healthy homeostasis.

**Fig. 2.**
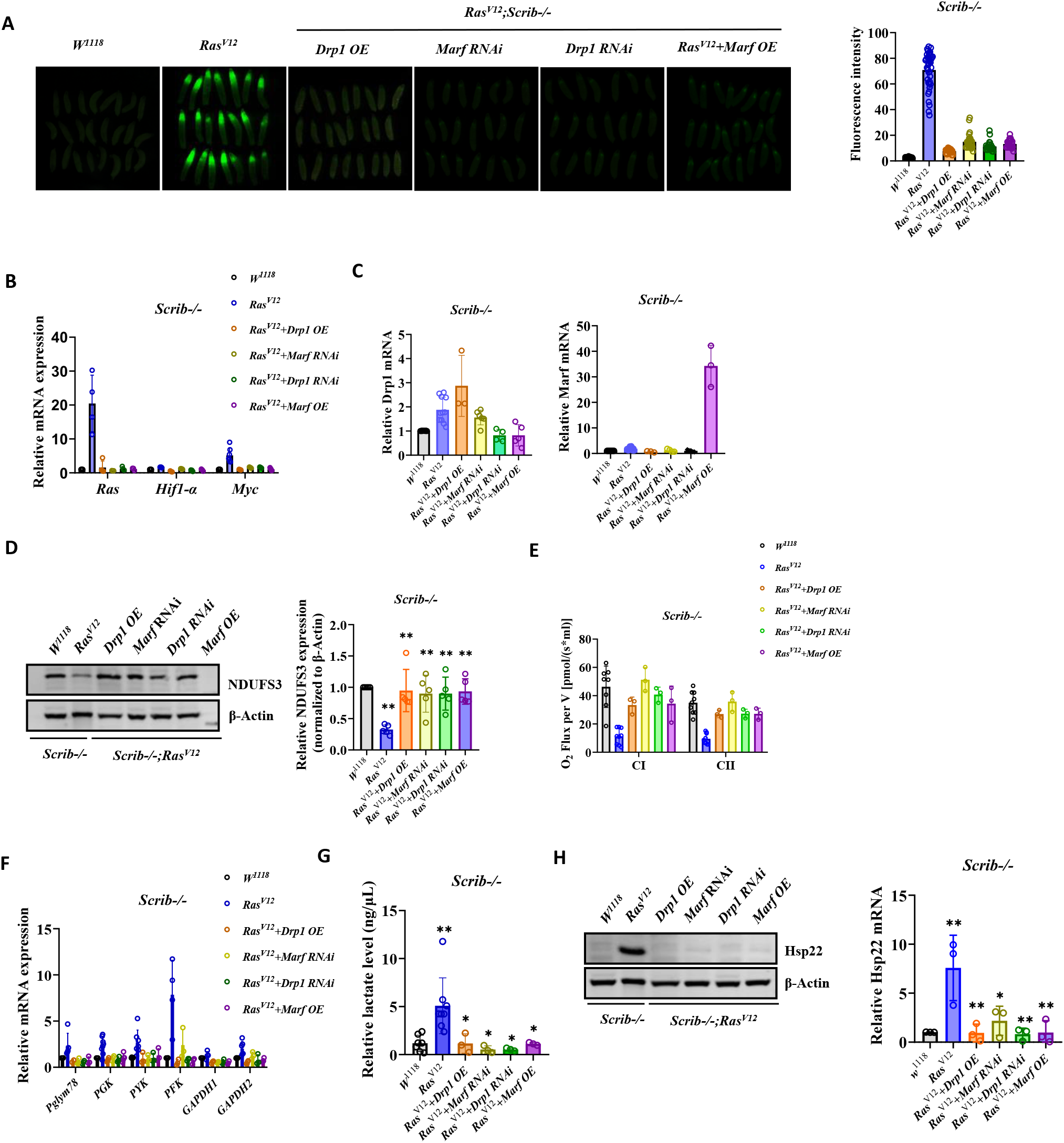
Tumor *Drosophila* model could be rescued by arbitrary modulation of mitochondrial dynamics with restored Hsp22 expression. **(A)** Arbitrarily modulated mitochondrial dynamics impact on *RasV12* tumor flies presented by GFP fluorescence. (**B**) Relative mRNA expression of Ras, Hif1-α and Myc in each group. (**C**) Relative Drp1 and Marf mRNA expression in each group. (**D**) NDUFS3 protein expression tested by WB in each group. (**E**) Quantification of mitochondrial CI and CII function tested in each group. (**F**) Glycolytic marker mRNA expression tested by RT-PCR in each group. (**G**) Relative lactate level tested separately in each group.. (**H**) Hsp22 protein expression tested separately in each group. All values are means ± SD of at least three independent experiments. Student’s t test (unpaired); *p< 0.05, **p< 0.01, **p< 0.001.

### 3 PD Drosophila model reprogrammed metabolism by modulation of mitochondrial dynamics with the restored Hsp22 expression

Loss of mitochondrial protein kinase PINK1 causes mitochondrial functional defects and degeneration of dopaminergic (DA) neurons which leads to PD development. As previous studies revealed that overexpression of Drp1 or downregulation of Marf rescued PINK1 mutant flies, we observed the same improved abnormal wing postures and motor actives (Figure 3A and B) with the increased tyrosine hydroxylase (TH) protein expression (Figure 3C) and rescued mitochondrial morphology tested by TEM (Figure 3D). Consistent with the restored mitochondrial morphology, mitochondrial ETC CI and CII function increased significantly and the increased NDUFS3 protein expression in Drp1 OE and Marf RNAi interfered PINK1B9 flies (Figure 3E and 3F). To our surprise, we found the dramatically decreased glycolysis capacity detected by RT-PCR and the highly induced lactate level which were rescued by mitochondrial dynamics interference (Figure 3G and 3H). More surprisingly, we found this improvement was associated with restored Hsp22 expression (Figure 3I) which further verified our hypothesis that Hsp22 is the key sensor in metabolism homeostasis.

**Fig. 3.**
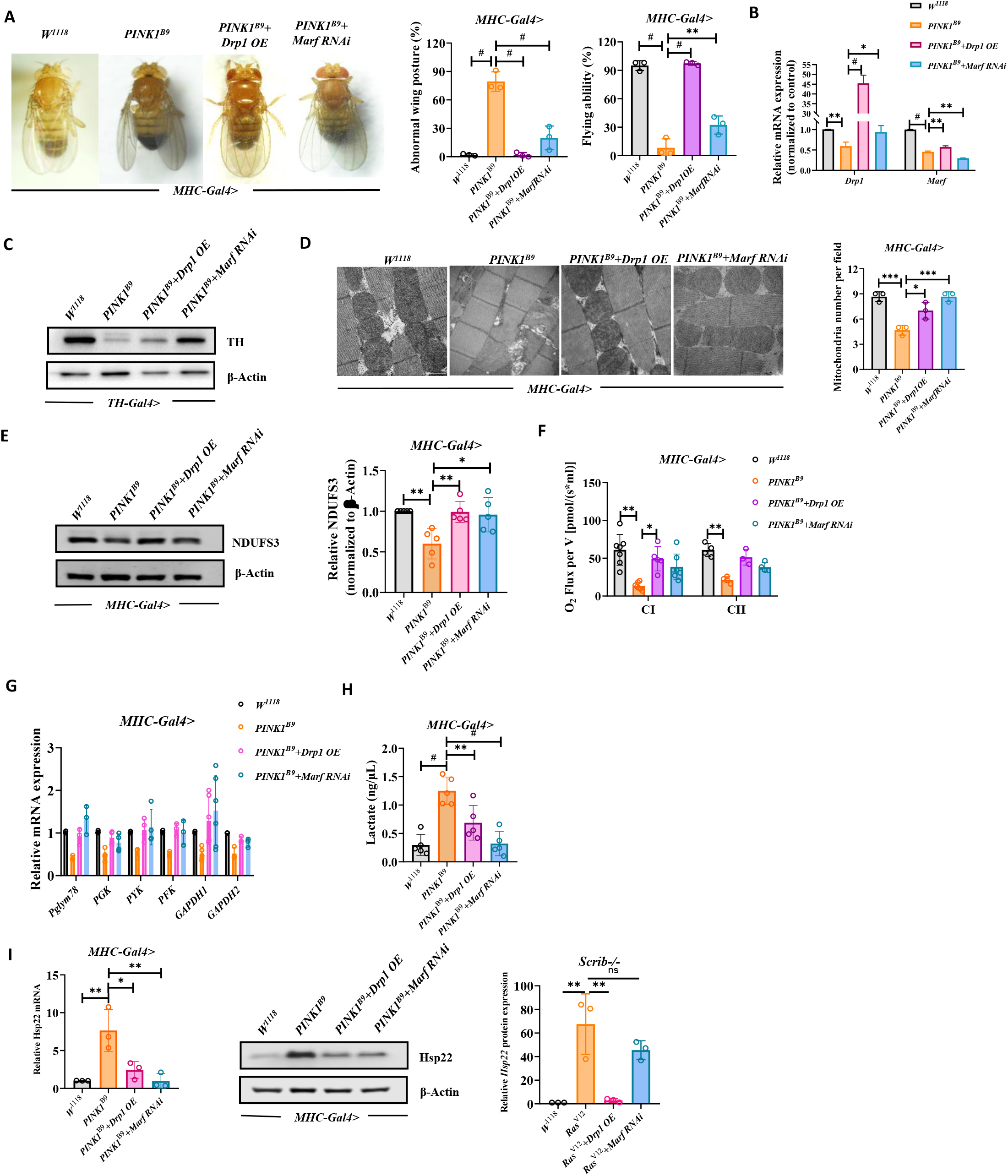
PD *Drosophila* model could be rescued by modulation of mitochondrial dynamics with the restored Hsp22 expression. (**A**) Mitochondrial fission trended modulation effect on PINK1B9 flies’ motor activity. (**B**) Relative *Marf* and *Drp1* mRNA expression tested by RT-PCR. (**C**) Tyrosine hydroxylase (TH) protein expression tested by WB separately in each group. (**D**) Mitochondrial morphology changes tested by TEM on mitochondrial fission trended modulation onPINK1B9 flies. (**E**) NDUFS3 protein expression tested by WB separately in each group. (**F**) Quantification of mitochondrial CI and CII function tested in thorax tissue. (**G**) Relative glycolytic marker of *MHC-Gal4* driven mitochondrial dynamics modulated mRNA expression. (**H**) Relative lactate level tested separately in each group. **(I)** Hsp22mRNA and protein expression tested separately in each group. All values are means ± SD of at least three independent experiments. Student’s t test (unpaired); *p< 0.05, **p< 0.01, **p< 0.001.

### 4 Overexpression of Hsp22 (Hsp22 OE) could reverse tumor models’ pathogenic phenotypes and metabolism

Hsp22 was induced in mitochondrial dynamics perturbation induced metabolism disturbed models as we described previously. We were wondering whether direct modulation of Hsp22 would rescue tumor pathogenic phenotypes and correct the pathogenic metabolism. Unexpectedly, we observed Hsp22 OE suppressed RasV12; Scrib-/- tumor flies’ pathogenesis revealed by the decreased in the clones marked with GFP fluorescence (Figure 4A). Furthermore, RT-PCR was conducted to confirm the suppression effect of Hsp22 OE on tumor invasion with decreased Ras, Myc and Hif1-α mRNA levels expression (Figure 4B). As expected, we observed the interfered mitochondrial dynamics alteration with the reduced Drp1 and Marf mRNA expression in RasV12; Scrib-/- flies (Figure 4C). In order to confirm whether Hsp22 OE could reverse the pathogenic metabolism in RasV12; Scrib-/- tumor flies, we tested the important subunit of CI in ECT which was confirmed by increased NDUFS3 expression by immunoblot as well (Figure 4D). More convincingly, we further observed that Hsp22 OE could directly increase CI and CII function compared with RasV12; Scrib-/- tumor flies tested by O2K equipment (Figure 4E). Interestingly, we found that PFK, GAPDH1, GAPDH2, PGK, Pglym78 and PYK mRNA in glycolysis pathway decreased significantly in Hsp22 OE RasV12; Scrib-/- tumor flies (Figure 4F) with the decreased lactate level (Figure 4G) indicating Hsp22 plays a very important role in reprogramming the pathogenic metabolism in RasV12; Scrib-/- tumor flies. The expression of Hsp22 in RasV12; Scrib-/- tumor flies were confirmed by WB (Figure 4H).

**Fig. 4.**
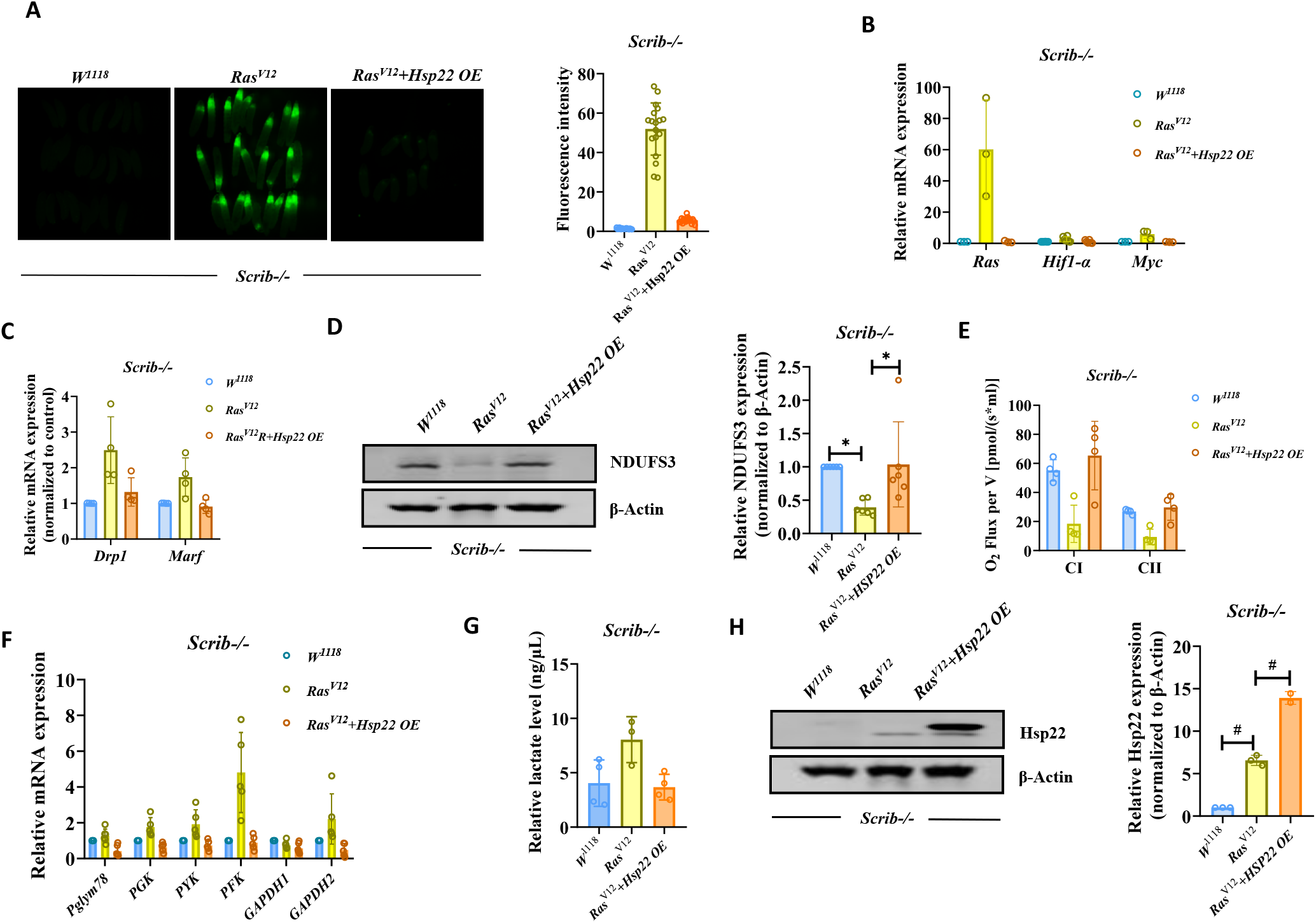
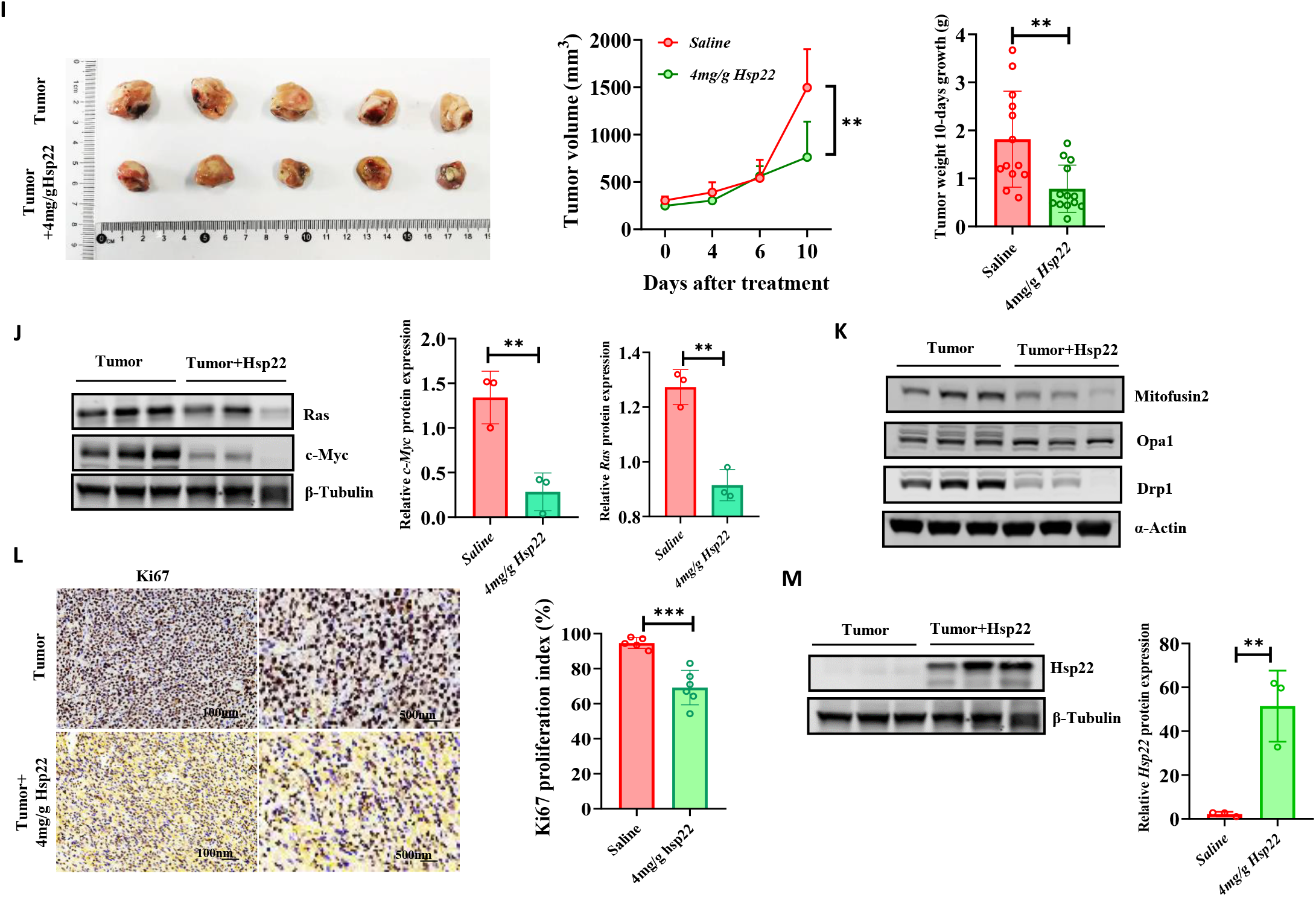
Overexpression of Hsp22 restores tumor models’ mitochondrial dynamic equilibrium and reverses pathogenic phenotypes and metabolism. **(A)** *Hsp22 OE* impacts on *RasV12* tumor flies presented by *GFP* fluorescence. (**B**) Relative mRNA expression of *Ras, Hif1-α* and *Myc* in each group. (**C**) Relative *Drp1* and *Marf* mRNA expression in each group. (**D**) *NDUFS3* protein expression tested by WB in each group. (**E**) Quantification of mitochondrial CI and CII function tested in each group. (**F**) Glycolytic marker mRNA expression tested by RT-PCR in each group. (**G**) Relative lactate level tested separately in each group.. (**H**) Hsp22 protein expression tested separately in each group. All values are means ± SD of at least three independent experiments. Student’s t test (unpaired); *p< 0.05, **p< 0.01, **p< 0.001. (**I**) In vivo monitoring of tumor formation of *H22* liver tumor with *Hsp22* supplement. (**J**) *Ras* and *c-Myc* protein expression tested by WB in each group. (**K**) *Mitofusion2, Opa1* and *Drp1* protein expression tested by WB in each group. (**L**) *Ki-67* staining of liver sections with 10-days *Hsp22* treatment. (**M**) *Hsp22* protein expression tested by WB in each group. All values are means ± SD of at least three independent experiments. Student’s t test (unpaired); *p< 0.05, **p< 0.01, **p< 0.001.

To test whether Hsp22 in the common corrector in tumor models, we generated HCC which is the most common liver tumor. Where introduced to confirm whether in vitro treatment of Hsp22 supplement would suppress tumor pathogenic process. In agreement with the results of previous studies, after 8 weeks, all of the mice injected with H22 cells bore tumors in the liver. As previous studies reported, the mice were treated with Hsp22 supplement (4mg/g) day and were sacrificed every 4 days for tumor parameters collection. Surprisingly, we observed the suppressed tumor growth by tumor volume detection especially after 10 days of Hsp22 treatment which was consistent with the tumor weight measured on the tenth day of Hsp22 treatment (Figure 4I). And we confirmed the decreased tumor exacerbation with the decreased Ras and c-Myc protein expression in tumor tissue (Figure 4J) with the rebalanced mitochondrial dynamics were tested by Mitofusin2, Opa1 and Drp1 protein expression in tumor tissue (Figure 4K). Since proliferation-associated nuclear antigen (Ki-67) Ki-67 is a tumor antigen that’s found in growing, dividing cells but is absent in the resting phase of cell growth, we stained the liver tumor tissue which presented the decreased Ki-67 expression in Hsp22 injected tumor mice compared with the saline treated tumor mice (Figure 4L). Hsp22 expression restoration was accompanied by the re-balanced mitochondrial dynamics (Figure 4M).

### 5 Overexpression of Hsp22 could reverse mitochondrial dynamics mediated PINK1B9 PD’s models’ pathogenic phenotypes and metabolism

The reversed metabolism was found in mitochondrial fission trended interference on PINK1B9 PD flies including improved glycolysis capacity (Figure 3) and elevated mitochondrial function which is consistent with previous studies. To test whether direct interfering PINK1B9 PD flies with Hsp22 OE could rescue the metabolic perturbation like the rescued tumor models. We therefore investigated whether overexpression of Hsp22 could influence PINK1B9 PD flies’ pathogenic phenotypes with abnormal wing posture and flying rates from which we found that Hsp22 OE significantly decreased PINK1B9 flies abnormal wing posture and increased flying ability (Figure 5A). To test whether Hsp22 interference would improve PINK1B9 flies mitochondrial morphology, we analyzed the ultrastructure of mitochondria using TEM. IFMs from control flies showed regular organization of myofibrils and densely packed mitochondria with intact cristae while PINK1 mutant displayed a heterogeneous population of mitochondria with the majority having significantly enlarged sizes and mild or severe disruption of their cristae structure, when compared to control mitochondria (Figure 5B). MHC-Gal4 driven Hsp22 OE significantly reduced the fraction of severely impaired mitochondria and increased the fraction of mitochondria with healthy cristae structure with the elevated Drp1 mRNA expression and decreased Marf mRNA levels (Figure 5C). Taken together, these observations suggest that overexpression of Hsp22 protects against the phenotypic consequences of PINK1 mutation. The mitochondrial dysfunction in PINK1B9 flies is shown by a loss of mitochondrial proteins and decreased respiration. By overexpressing Hsp22 in PINK1B9 flies, we found the dramatic increase of CI and CII function of ETC while we also observed a recovery of the mitochondrial protein content, indirectly assessed through the analysis of the levels of CI subunit, NDUFS3 (Figure 5D and E). More importantly, we were wondering whether the improvement of mentalism from Hsp22 overexpression would alter glycolysis metabolism too since the decreased glycolysis was another hallmark reported recently in PD patients which was also proved in our previous findings. We noted that PFK, GAPDH1, GAPDH2, PGK, Pglym78 and PYK mRNA in glycolysis pathway increased significantly in Hsp22 OE rescued PINK1B9 flies accompanied by the decreased lactate generation (Figure 5F and G). We detected the increase in the Hsp22 protein level in PINK1B9 flies in thorax tissue that was reversed upon the overexpression of Hsp22 which was opposite to our expectation (Figure 5H). This result indicates that overexpression of Hsp22 supplemented the protective induction of Hsp22 in PINK1B9 PD flies and reversed metabolism reprogramming in PINK1B9 PD flies.

**Figure 5.**
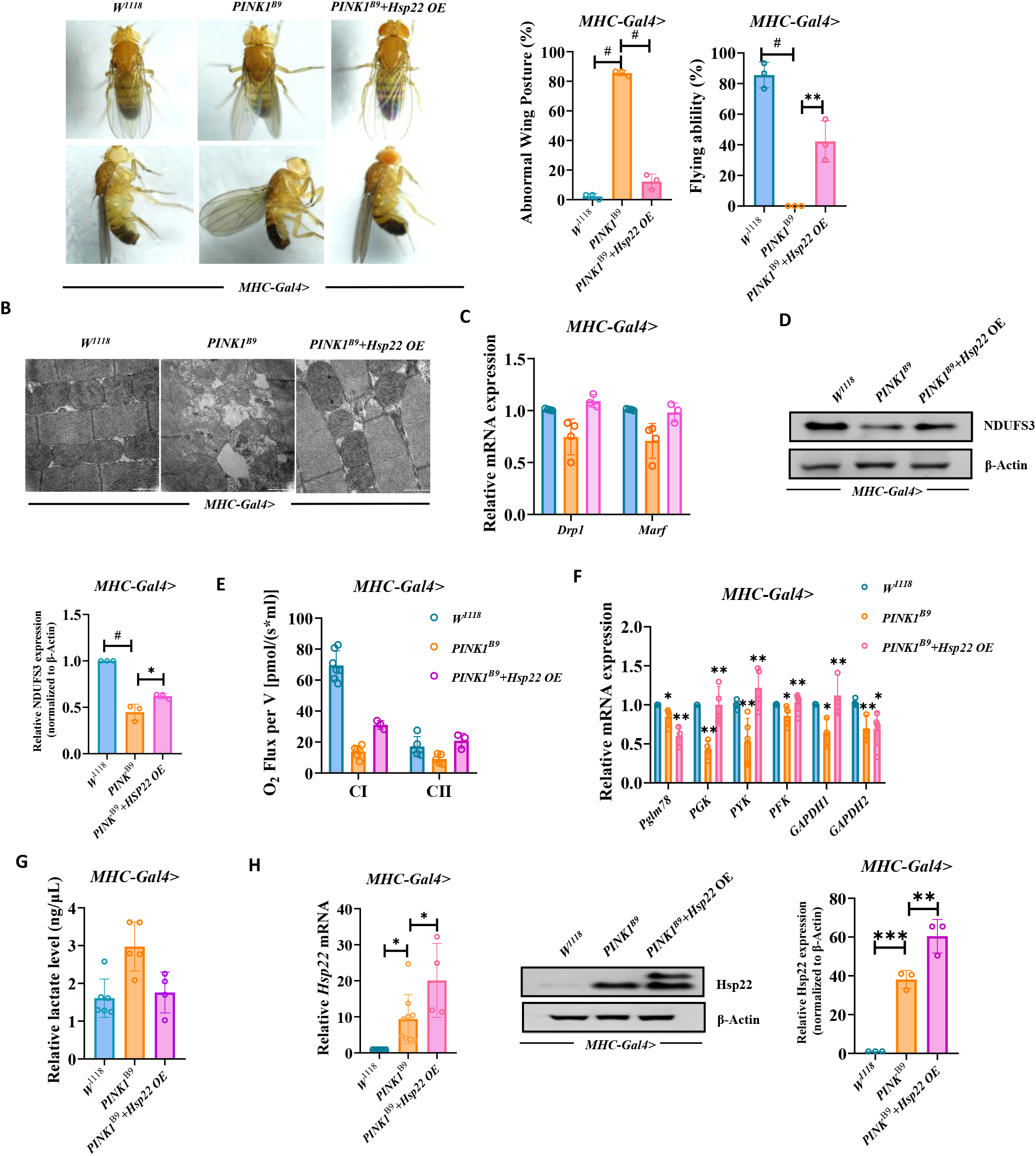
Overexpression of Hsp22 could reverse PINK1B9 PD’s mitochondrial dynamic equilibrium, pathogenic phenotypes and metabolism. (**A***) Hsp22 OE* impacts on *PINK1B9* flies’ motor activity. (**B**) Mitochondrial morphology changes tested by TEM on mitochondrial *Hsp22 OE* interference on *PINK1B9* flies. (**C**) Relative *Marf* and *Drp1* mRNA expression tested by RT-PCR. (**D**) *NDUFS3* protein expression tested by WB separately in each group. (**E**) Quantification of mitochondrial CI and CII function tested in thorax tissue. (**F**) Relative glycolytic marker of Hsp22 OE modulated thorax mRNA expression. (**G**) Relative lactate level tested separately in each group. (**H**) Hsp22mRNA and protein expression tested separately in each group. All values are means ± SD of at least three independent experiments. Student’s t test (unpaired); *p< 0.05, **p< 0.01, **p< 0.001.

### 6 ‘Neutralizing’ tumor pathogenic metabolism with PD induction factor accompanied by mitochondrial dynamic associated Hsp22 modulation

Since we observed the severely induced Hsp22 expression in mitochondrial dynamics perturbated models including tumor and PD whose pathogenesis were regarded as biochemically opposite. From our previous demonstrations, we found the potential opposite mechanism between tumor and PD. From the same initiation, mitochondrial dynamics disorder induced highly balanced dynamics in tumor accompanied by the high glycolysis capacity which is totally opposite to PD whose mitochondrial dynamics were impaired significantly accompanied by the significantly decreased glycolysis capacity. We were wondering whether the pathogenic symptoms and metabolism could be reversed by the disease induction factors in these two diseases. To address this, the environmental factor, which is well studied in causing parkinsonism, 1-methyl-4-phenyl-1, 2, 3, 6-tetrahydropyridine (MPTP) was introduced to our investigation. Surprisingly, we found that 24mM MPTP treated RasV12; Scrib-/- tumor flies for 5 days presented the significant suppression on tumor growth compared with the untreated controls tested by clones marked with GFP fluorescence (Figure 6A). To understand whether this suppressive effect on RasV12; Scrib-/- tumor flies could further impact on the cellular pathogenic metabolism, we moved on to test the signature genes of glycolysis pathway which revealed that PFK, GAPDH1, GAPDH2, PGK, Pglym78 and PYK mRNA levels significantly decreased and resulted in the decrease of lactate level in MTPT treated ones compared with the untreated tumor flies (Figure 6E). To refine the mitochondrial metabolic reprogramming involved in the protective effect of MPTP in RasV12; Scrib-/- tumor flies, we tested mitochondrial function by ETC CI and CII O2 flux from which we observed a significant increase in MPTP treated Scrib-/- tumor flies’ tissue with increased NDUFS3 protein level compared with the tumor controls (Figure 6C). The relatively restored cellular metabolism in MPTP treated RasV12; Scrib-/- tumor flies might be due to the restored Hsp22 associated mitochondrial dynamics. To address this, we tested the expression of Drp1, Marf and Hsp22 mRNA expression from which we observed the significant decrease of Drp1 and Marf mRNA expression (Figure 6G) indicating the pathogenic highly induced metabolism was suppressed with the restored lowered Hsp22 mRNA expression (Figure 7E).

**Fig. 6.**
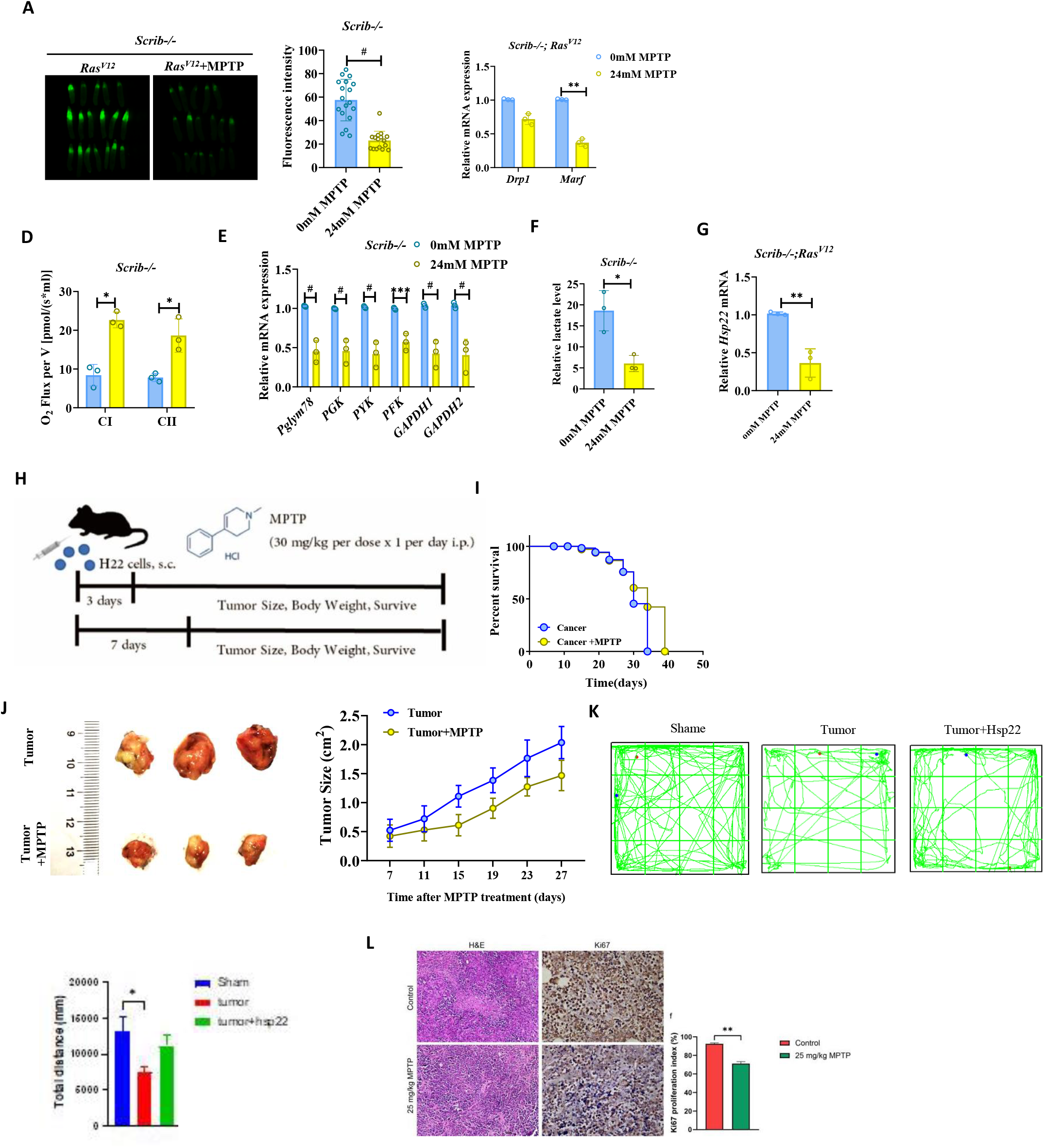
Tumor models could be rescued by PD induction factor accompanied with Hsp22 modulation. (**A**) *MTPT* treatment impacts on *RasV12* tumor flies presented by *GFP* fluorescence. (**B**) Relative *Marf* and *Drp1* mRNA expression tested by RT-PCR.. (**C**) *NDUFS3* protein expression tested by WB in each group. (**D**) Quantification of mitochondrial CI and CII function tested in each group. (**E**) Glycolytic marker mRNA expression tested by RT-PCR in each group. (**F**) Relative lactate level tested in each group. (**G**) *Hsp22* mRNA expression tested separately in each group. (**H**) *MPTP* treating strategy on *H22* liver tumor mouse model. (**I**) Monitored survival of *H22* liver tumor mice with *MPTP* treatment. (**J**) Monitored *H22* tumor size with *MPTP* treatment. (**K**) Open-field tests on *MPTP* treated *H22* tumor mice. (**K**) *Ki-67* staining of liver sections with 10-days *MPTP* treatment.

To further confirm whether PD induction factor would restore the pathogenic metabolism in mammal tumor models, here we treated liver tumor mice with the same PD induction factor, MPTP followed by the construction strategy shown in Figure 7F. As we speculated that with MPTP PD induction factor treatment liver tumor mice resulted in prolonged survival (Figure 7G). Consistent with the protective role in survival assay, we found that the liver tumor size development was suppressed in MPTP treated tumor mice under the continuous monitoring in tumor size measurements (Figure 7H). Also, we found the significantly decreased tumor weight in MPTP treated liver tumor mice detected at the ending time of monitoring (Figure 7I). To further determine whether the suppressive role in liver tumor invasion of MPTP PD induction factor was involved in mitochondrial dynamics alteration, we tested Drp1, Marf and Opa1 levels tested by immunoblots from which we found the significant decreased protein expression levels of Drp1, Marf and Opa1 separately in MPTP treated liver tumor mice (Figure 7J).

## Discussion

Classically, mitochondria play an essential role in cellular bioenergetics and maintaining cellular homeostasis with fission and fusion dynamic equilibrium that is mediated by notably Marf which promotes fusion, and Drp1 which promotes fission [9]. Under healthy status, the dynamic mitochondrial fission and fusion cycle is proposed to balance two competing processes: elimination of damage by fission and compensation of damage by fusion [10]. However, failure of mitochondrial dynamics regulation results in disrupted cellular metabolism which is widely thought to lead to numerous diseases including tumor, PD and even the natural aging since mitochondria-associated metabolic activity is believed to decrease gradually during aging [11]. Here we provide the evidence for mitochondrial dynamics regulation induces cellular metabolic reprogramming including the reprogrammed glycolysis capacity and mitochondrial function which can be presented by tumor and PD as the specific mitochondrial dynamic defects’ models.

We hypothesize that mitochondrial dynamic balance is like a seesaw in oscillation, and changes in fission or fusion extent will cause the mitochondrial dynamics to gradually become imbalanced, the oscillation frequency will become lower and lower, and finally will be halted due to the extremely imbalanced mitochondrial dynamics and even may end up in apoptosis. When mitochondrial fission and fusion extent increase at the same time which leads to the new rebalanced status represented by tumor. While the frequency of seesaw oscillation increases in mitochondrial dynamics which leads to the continuous tumor invasion. This seesaw model might explain why tumor and PD is barely seen in one single patient. And also proved by PD induction factor, no matter genetically interference (PINK1 and Parkin) in tumor flies or MPTP treatment in liver mouse model both showed the suppressed tumor invasion and the reversed pathogenic metabolism indicating tumor and PD might share the oppositely developed pathogenic metabolism which explains the “neutralizing” effect in tumor with PD induction treatment.

Any perturbations in mitochondrial fission or fusion, including tumor and PD, will induce the mitochondrial dynamic repairing factor Hsp22 which plays the protective role in correcting abnormal mitochondrial dynamics and metabolic reprogramming. Among various Hsps, we have demonstrated that the expression of Hsp22 is strongly upregulated in mitochondrial dynamics disturbed models including tumor and PD. Hence, Hsp22 associated with the altered mitochondrial function and glycolysis capacity. We have also demonstrated that genetically upregulation of Hsp22 could suppressed RasV12 flies’ tumor invasion and improve cellular metabolism and moreover, we also proved that exogenous injection of Hsp22 protein on mouse hepatocellular carcinoma model could reverse the pathogenic phenotypes and metabolism. In line with PINK1B9 PD flies’ mitochondrial dynamics defects, we proved that Hsp22 OE improved PINK1B9 PD flies’ neurodegenerative phenotypes and the improved metabolism which also highly demonstrates the key sensor and balancer role of Hsp22 in mitochondrial dynamics associated defects.

In conclusion, our study reveals a key role of Hsp22 in regulating cellular metabolism via mitochondrial dynamics. We suggest that Hsp22 plays partially as the key common sensor and balancer in mitochondrial dynamics dysregulation-initiated diseases proved in our study indicating Hsp22 may be the potential therapeutic target for mitochondrial dynamic dysregulation mediated metabolic reprogrammed diseases including tumor, PD and managing a well natural aging.

## Supporting information

Seesaw movie

## Supplementary Material

## Supp

### Material and metochds

#### Fly Strains

Flies were maintained on standard food at 25°C with 70% humidity. All strains were obtained from the Bloomington Drosophila Stock Center, from the VDRC, or as gifts from colleagues. All fly crosses were carried out at 25°C with standard laboratory conditions unless noted otherwise. The following fly strains were used in this study: w1118 and PINK1^B9^ were obtained from Bloomington Stock Center (stock ID: 5905 and 34749); Mhc-GAL4 were donated by Fuzhou university; UAS-Marf-flag were donated by Leo Pallanck; Drp1 DNDK1161D were donated by fuzhou university; MarfRNAi were obtained from Bloomington Stock Center (stock ID: 31157); UAS-Drp1 were donated by Mel Ferny; UAS-Drp1-HA were donated by Ming Guo; PINK1 RNAi and Parkin OE were donated by The state key lab of medical genetics; Parkin RNAi were obtained from Bloomington Stock Center (stock ID: 38333); PINK1OE were obtained from Bloomington Stock Center (stock ID: 51648); Drp1 RNAi were obtained from Bloomington Stock Center (stock ID: 27682); Scrib-GAL4 and RasV12 were donated by Lei Xue, Professor of Tongji University in Shanghai.

#### Drosophila flying rate and abnormal wing posture

If the wings overlap completely and are horizontal to the body, it is normal; if the wings are vertical, forked or drooping, it is abnormal, and the rate of wing anomaly is calculated. Statistical anomaly rate. The glass tube was tapped with the same frequency to observe the jumping behavior of the target flies, and the number of jumping and flying flies was recorded. Finally, the flight rate was counted. It is important to exclude the jumps caused by the vibration of the glass tube.

Total RNA was isolated from thorax tissue of drosophila or larvas by using Trizol Reagent (Invitrogen, Carlsbad, CA, USA).^25^ cDNA was generated using the Prime Script™ RT reagent Kit with gDNA Eraser (TAKARA RR047A, Japan).^26,27^ Random primers were added as primer for reverse transcription. According to the manufacturer’s instruction. Further Q-PCR was performed using specific primers and SYBR Green QPCR mix on ABI 7500 Real Time PCR system, via amplification steps as followed:50L 30sec, 95L 10 min, 95L 15sec, then 60L 1min.The Ct Values were then calculated according to the flue curves for each reaction. 18s was used as a control. Sequences of the primers used in this study were as follows: 18s, 5’-TCTAGCAATATGAGATTGAGCAATAAG -3’ and 5’-AATACACGTTGATACTTTCATTGTAGC -3’, Pfk, 5’-ATCGTTGGATTGGTTGGCTC -3’ and 5’-TGCCTCGATGATGCGATGTAG -3’, Pgk, 5’-TGATGCGCGTCGACTTCAAT-3’ and 5’-TTGATACTATCCAAGGCAGCGA-3’, Pyk, 5’-CATCCGCATTGTCACCGTC-3’ and 5’-AGTAATCAACGCGACGCCC-3’, Pglym78, 5’-AGCATTTAGACAACCTTTCTGAGGA-3’ and 5’-GTTCTCGTCCAGCTCGTAGAC-3’, GAPDH1, 5’-AGCCAAAACTATCGTACAAACCC-3’ and 5’-CATCCTCAATTGCGCCCCTT-3’, GAPDH2, 5’-ACACTACCCACCCACACTCT-3’ and 5’-GTGATGCCATTTTCAGGCCG-3’, Ras, 5’-TGGTTATCGATGGAGAGACCTG -3’ and 5’-TCCGCATATACTGATCCCGC-3’, P53, 5’-TGAGCCTTTGACGGCCAATA-3’ and 5’-AATTCCCTTGCCCTGAGCAT-3’, Myc, 5’-GAAGAACAACAGCAACGGCAT-3’ and 5’-TTTCCTCATGGAATCTGAGGGG-3’, Hif1-α, TGACGATTCCGAAGCAATGA and ACGTGGAGGTATGACAGTGC, Marf, 5’-GTATTGGACACAGCGTATCGAC-3’ and 5’-AGCTGCTCGGGCGTAATG-3’, Drp1, 5’- AGGCCCTAATTCCGGTCATA-3’ and 5’-CGGAGCTCTGACTGCCTAGA-3’, Parkin, 5’-ACGACAATAGAGCAATGTGACT-3’ and 5’-CGCTAAGCGAAGGTTCCTC-3’, Pink1, 5’-ACGTGATATACACGCCAACAT-3’ and 5’-CTGCTGGGGCGTTGTCC-3’, Hsp22, 5’-GAGGTGACCATCGAGCAGAC-3’ and 5’-TTCTACTGACTGGCGGCTTT-3’, Hsp23, 5’-AGCGAACTGGTGGTCAAAG-3’ and 5’-GGACAAAGTGACGAGTGATGAA-3’, Hsp26, 5’-GGAGCGCATCATTCAAATTCAG-3’ and 5’-TGGCTCCTTTACTTGTCCTTG-3’, Hsp27, 5’-CTGGGTCGTCGTCGTTATTC-3’ and 5’-CACACCTGGAAGCCATCTT-3’, Hsp68, 5’-CGCAATCAACTGGAGACCTATT-3’ and 5’-TGCTGTCCAACCACTTCATC-3’, Hsp83, 5’-GAGGATGAGGAGCTGAACAAG-3’ and 5’-CAGTCGTTGGTCAGGGATTT-3’.

#### Electron microscopy

The target Drosophila was anesthetized with ether and rapidly cut off the head, then put into 3% glutaraldehyde fixing solution, fixed at 4°C for 24h. Then washed with 0.1M phosphate buffer for 3 times, 1% eucalyptus fixed for 2h, then dehydrated with ethanol - acetone. The acetone-embedding agent permeated for more than 2h, and then after embedding-polymerization-block repair steps, the sections were cut with (LAIKA UC7) ultra-thin slices, uranium acetate - lead citrate double staining, and finally observed with electron microscope (Hitachi H-7650).

#### Measurements of fly’s respiration

Mitochondrial respiration was assayed at 37 °C by high-resolution respirometry using an OROBOROS Oxygraph. DatLab software package (OROBOROS, Innsbruck, Austria) was used for the data acquisition (2-s-time intervals) and analysis. Respiration was assayed by homogenising five flies using a pestle in 100μl MiR05 respiration buffer (20mM HEPES, 10mM KH2PO4, 110mM sucrose, 20mM taurine, 60mM K-lactobionate, 0.5mM EGTA, 3mM MgCl2, 1 g/L fatty acid-free BSA). complex I activity was examined in MiR05 respiration buffer after titration with 5mM pyruvate, 2mM malate, 10mM glutamate and 5mM ADP. 10μM Cytochrome c was used to detect mitochondrial membrane integrity. Complex II was determined after adding 0.5μM rotenone and 10mM succinate. Residual oxygen consumption was evaluated after the addition of 2.5 μM.

#### Lactate quantification

Amount of lactate in tissues of flies (n=20) was estimated using Lactate Assay Kit (Sigma, Catalog Number MAK064) following manufacturer’s instructions.

#### Immunofluorescence methodology and intensity analysis methods

The RasV12; scribe-/- tumor Drosophila containing GFP-tagged FLP/FRT recombinant system may observe significant fluorescence under Nikon SMZ18 body fluorescence microscope, and then its fluorescence intensity was calculated using the NIS-Elements Imaging Software, which comes with the instrument’s own analysis software.

#### The modeling method for mouse liver tumor

C57BL/6J mice were purchased from Hunan Slake Jingda Experimental Animal Co. 6∼8 weeks old C57BL/6J male mice (bodyweight, 23±3g) were intraperitoneally injected H22 cells. 20 mice were randomly selected to receive a subcutaneous injection of 0.2ml ascites cells (107/ml) in the groin for about a week. The tumor size was visually measured at 0.8mm, and the drug was administered.

#### Preparation of Hsp22 protein lyophilizer

Gene sequence synthesis and insertion of the hsp22 protein into the pET28a vector successfully construct an expression plasmid. Introduce the expression vector into E. coli and pick out single colonies in 10 ml of LB medium (canna-resistant) at 37°C, 250 rpm, overnight. Transfer 1% overnight bacteria in 1000 ml of LB medium (Canadiene-resistant) at 37°C at 250 rpm for 3 hours, then add 0.5 mM IPTG at 20°C for 12 hours of induction. Collect 1000 ml of E. coli culture expressing the target protein by centrifugation, sonicate (300 W, sonication for 10 seconds, an interval for 10 seconds, times 30) in 100 ml of bacterial breaking buffer (50 mM PBS, pH 7.4, 0.15 M NaCl), and collect the precipitate and supernatant after centrifugation, respectively. Take the supernatant after centrifugation, dissolve it with PBS and sample it on a pre-equilibrated NTA purification column (1 ml NTA medium, equilibrated with 10 times the volume of the column in NTA-0 buffer). After sample loading, wash the column with NTA-0 buffer for 10 column volumes. Wash the column sequentially with NTA-X buffer (50 mM PBS, pH 7.4, 0.15 M NaCl, X mM imidazole at different imidazole concentrations 10/20/50/100/200/500mM) by 1 column volume each. Step-by-step collection and electrophoresis of the different fractions, with the final mixture of the target protein’s fractions will be the components with PBS dialysis to remove salts and heteroproteins, the resulting protein liquid lyophilized into powder. Dissolve the protein lyophilizer in pure water to a concentration of about 20mg/ML and inject it into rats at a concentration of 4mg/kg.

#### Hsp22 treating method

The above-mentioned model mice with uniform tumor size were selected and randomized into groups: sham-operated group, group given hsp22 protein alone, tumor group, tumor group given hsp22 protein, six in each group. hsp22 protein administration group was given 4 mg/ml of purified protein intraperitoneally every day, and sham-operated group and tumor group alone were given an equal volume of saline.

#### Mouse liver tumor weight and volume determination method

From the first day of administration, the tumor size (length and width) was measured daily with a Vernier caliper, and after 10 days, the mice were killed, the tumor body was dissected out, the filter paper was drained of body fluid, weighed and recorded.

#### Mouse immunohistochemistry methods and statistical methods (Ki67)

Tumor tissues were fixed with 4% paraformaldehyde for 24h, dehydrated with ethanol solution at all levels, transparent treated with xylene, made into paraffin blocks, sectioned with a microtome, dewaxed and rehydrated, repaired with antigen, incubated with ki67 (abcam 1:1000) primary antibody at 4°C overnight, incubated with secondary antibody (goat anti-rabbit 1:2000), developed by DAB, dehydrated and sealed, and photographed under the microscope. The slides were photographed under a microscope (Leica) at 40×, and 6 fields of view were randomly selected for each slice, counted, and GraphPad Prism 8.0 was used for statistical results.

#### Drosophila tumor MPTP treating method

The promoter scrib-GAL4, which is selectively expressed in the larval eye disc and abdominal nerve cord of Drosophila larvae, was crossed with RasV12 Drosophila, which is fluorescently labeled with GFP, in normal culture medium. After 12 days of MPTP administration, larvae were picked and observed under a fluorescence microscope and the fluorescence intensity was calculated.

#### MPTP administration method in tumor mice

12 mice with similar body weight and age were randomly divided into sham-operated group and MPTP-treating group. The tumor group model was constructed as above, and when the tumor size was about 0.8mm, 12 mice with similartumor size were selected and randomly divided into tumor group and MPTP administration group. The MPTP administration group was given MPTP (25 mg/kg) intraperitoneally every day, and the sham-operated and tumor groups were given an equal volume of saline. The drug was administered continuously for one week.

#### MPTP tumor mouse survival analysis method

The tumor model and MPTP administration were the same as above, and the time of death of each group of mice was observed and recorded to calculate life span.

#### MPTP treated tumor volume analysis

The length (a) and width (b) of the tumor were measured by Vernier calipers, and the volume of the tumor in each group was calculated according to the formula v=a×b2 to draw the growth curve of the tumor.

#### Absenteeism experiment

Absentee field experimental animal behavior analysis system (provided by Shanghai XinSoft Information Technology Co., Ltd.), the mice into a clean, odorless absentee box, and timed for 5min, while the camera and timing, the experiment in a quiet environment. Clean the inside wall and bottom of the box to avoid the residual information of the last animal (such as animal’s urine and faeces, smell) affect the next test results. Replace the animals and continue the experiment.

The data were collected from the total distance, and the results were recorded by GraphPad Prism 8.0.

## References and Notes

1. Rugarli, E. I., and T. Langer. 2012. Mitochondrial quality control: a matter of life and death for neurons. The EMBO journal 31:1336–1349. doi 10.1038/emboj.2012.38

2. Kashatus, J. A., A. Nascimento, L. J. Myers, A. Sher, F. L. Byrne, K. L. Hoehn, C. M. Counter, and D. F. Kashatus. 2015. Erk2 phosphorylation of Drp1 promotes mitochondrial fission and MAPK-driven tumor growth. Molecular cell 57:537–551. doi 10.1016/j.molcel.2015.01.002

3. Civenni, G., R. Bosotti, A. Timpanaro, R. Vàzquez, J. Merulla, S. Pandit, S. Rossi, D. Albino, S. Allegrini, A. Mitra, S. N. Mapelli, L. Vierling, M. Giurdanella, M. Marchetti, A. Paganoni, A. Rinaldi, M. Losa, E. Mira-Catò, R. D’Antuono, D. Morone, K. Rezai, G. D’Ambrosio, L. Ouafik, S. Mackenzie, M. E. Riveiro, E. Cvitkovic, G. M. Carbone, and C. V. Catapano. 2019. Epigenetic Control of Mitochondrial Fission Enables Self-Renewal of Stem-like Tumor Cells in Human Prostate Cancer. Cell metabolism 30:303-318.e306. doi 10.1016/j.cmet.2019.05.004

4. McCoy, M. K., and M. R. Cookson. 2012. Mitochondrial quality control and dynamics in Parkinson’s disease. Antioxidants & redox signaling 16:869–882. doi 10.1089/ars.2011.4019

5. Taylor, R. P., and I. J. Benjamin. 2005. Small heat shock proteins: a new classification scheme in mammals. Journal of molecular and cellular cardiology 38:433–444. doi 10.1016/j.yjmcc.2004.12.014

6. Qiu, H., P. Lizano, L. Laure, X. Sui, E. Rashed, J. Y. Park, C. Hong, S. Gao, E. Holle, D. Morin, S. K. Dhar, T. Wagner, A. Berdeaux, B. Tian, S. F. Vatner, and C. Depre. 2011. H11 kinase/heat shock protein 22 deletion impairs both nuclear and mitochondrial functions of STAT3 and accelerates the transition into heart failure on cardiac overload. Circulation 124:406–415. doi 10.1161/circulationaha.110.013847

7. Bouhy, D., M. Juneja, I. Katona, A. Holmgren, B. Asselbergh, V. De Winter, T. Hochepied, S. Goossens, J. J. Haigh, C. Libert, C. Ceuterick-de Groote, J. Irobi, J. Weis, and V. Timmerman. 2018. A knock-in/knock-out mouse model of HSPB8-associated distal hereditary motor neuropathy and myopathy reveals toxic gain-of-function of mutant Hspb8. Acta neuropathologica 135:131–148. doi 10.1007/s00401-017-1756-0

8. Morrow G, Tanguay RM. rosophila melanogaster Hsp22: a mitochondrial small heat shock protein influencing the aging process. Front Genet. 2015 Mar 16; 6:1026. doi: 10.3389/fgene.2015.00103. PMID: 25852752; PMCID: PMC4360758.

9. van der Bliek, A. M., Q. Shen, and S. Kawajiri. 2013. Mechanisms of mitochondrial fission and fusion. Cold Spring Harbor perspectives in biology 5. doi 10.1101/cshperspect.a011072

10. Archer, S. L. 2013. Mitochondrial dynamics--mitochondrial fission and fusion in human diseases. The New England journal of medicine 369:2236–2251. doi 10.1056/NEJMra1215233

11. Corrado M, Scorrano L, Campello S. Mitochondrial dynamics in cancer and neurodegenerative and neuroinflammatory diseases. Int J Cell Biol. 2012; 2012:729290. doi:10.1155/2012/729290

