## Supplementary material for "Hsp22 is the key sensor and balancer in mitochondrial dynamic associated metabolic reprogramming": Supp1

### Slide 1
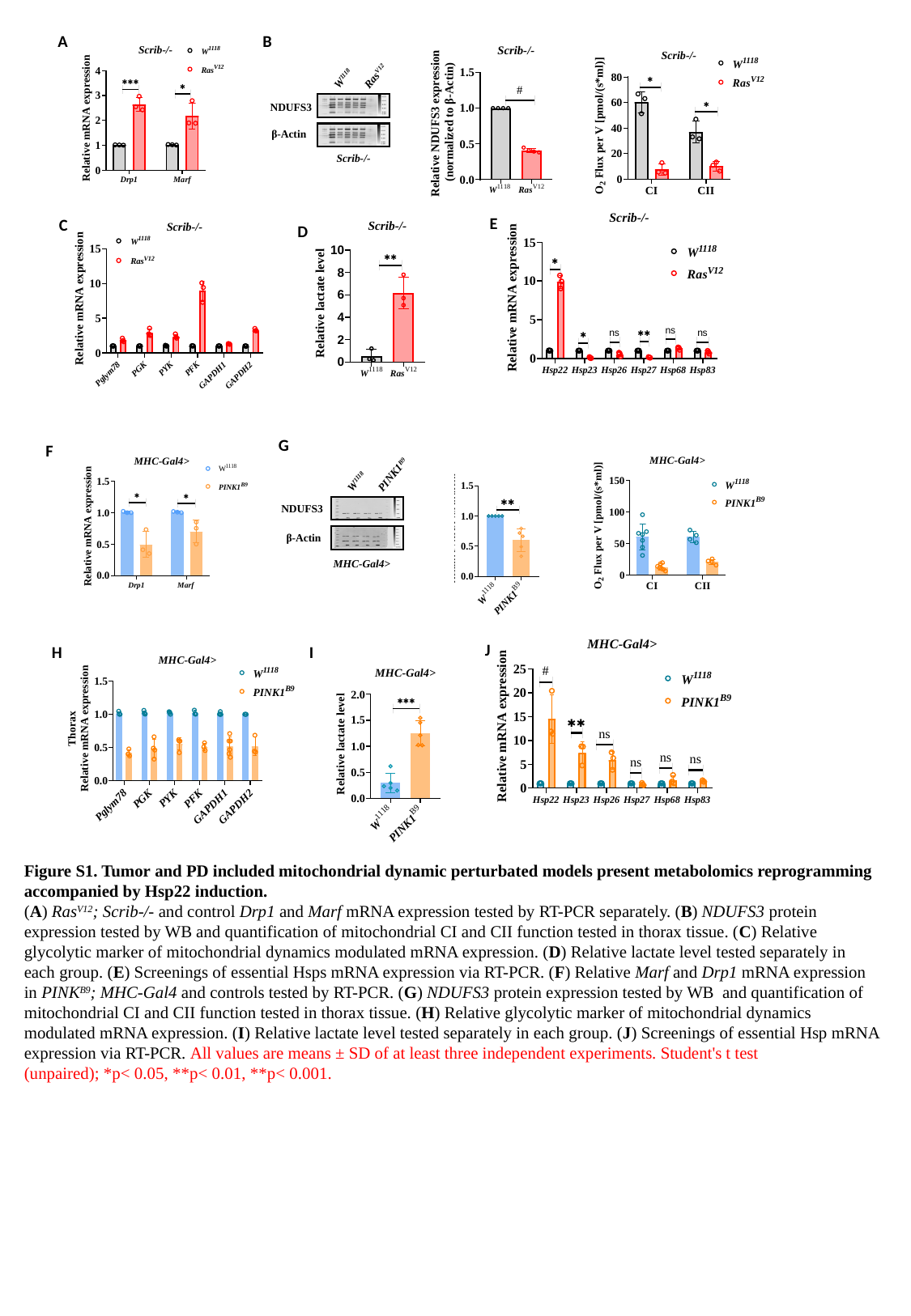

A
B
Ras^V12^
W^1118^
NDUFS3
β-Actin
Scrib-/-
E
C
D
G
F
PINK1^B9^
W^1118^
NDUFS3
β-Actin
MHC-Gal4>
J
I
H
Figure S1. Tumor and PD included mitochondrial dynamic perturbated models present metabolomics reprogramming accompanied by Hsp22 induction.
(A) Ras^V12^; Scrib-/- and control Drp1 and Marf mRNA expression tested by RT-PCR separately. (B) NDUFS3 protein expression tested by WB and quantification of mitochondrial CI and CII function tested in thorax tissue. (C) Relative glycolytic marker of mitochondrial dynamics modulated mRNA expression. (D) Relative lactate level tested separately in each group. (E) Screenings of essential Hsps mRNA expression via RT-PCR. (F) Relative Marf and Drp1 mRNA expression in PINK^B9^; MHC-Gal4 and controls tested by RT-PCR. (G) NDUFS3 protein expression tested by WB and quantification of mitochondrial CI and CII function tested in thorax tissue. (H) Relative glycolytic marker of mitochondrial dynamics modulated mRNA expression. (I) Relative lactate level tested separately in each group. (J) Screenings of essential Hsp mRNA expression via RT-PCR. All values are means ± SD of at least three independent experiments. Student's t test (unpaired); *p< 0.05, **p< 0.01, **p< 0.001.
