## Supplementary material for "Hsp22 is the key sensor and balancer in mitochondrial dynamic associated metabolic reprogramming": Supp2

### Slide 1
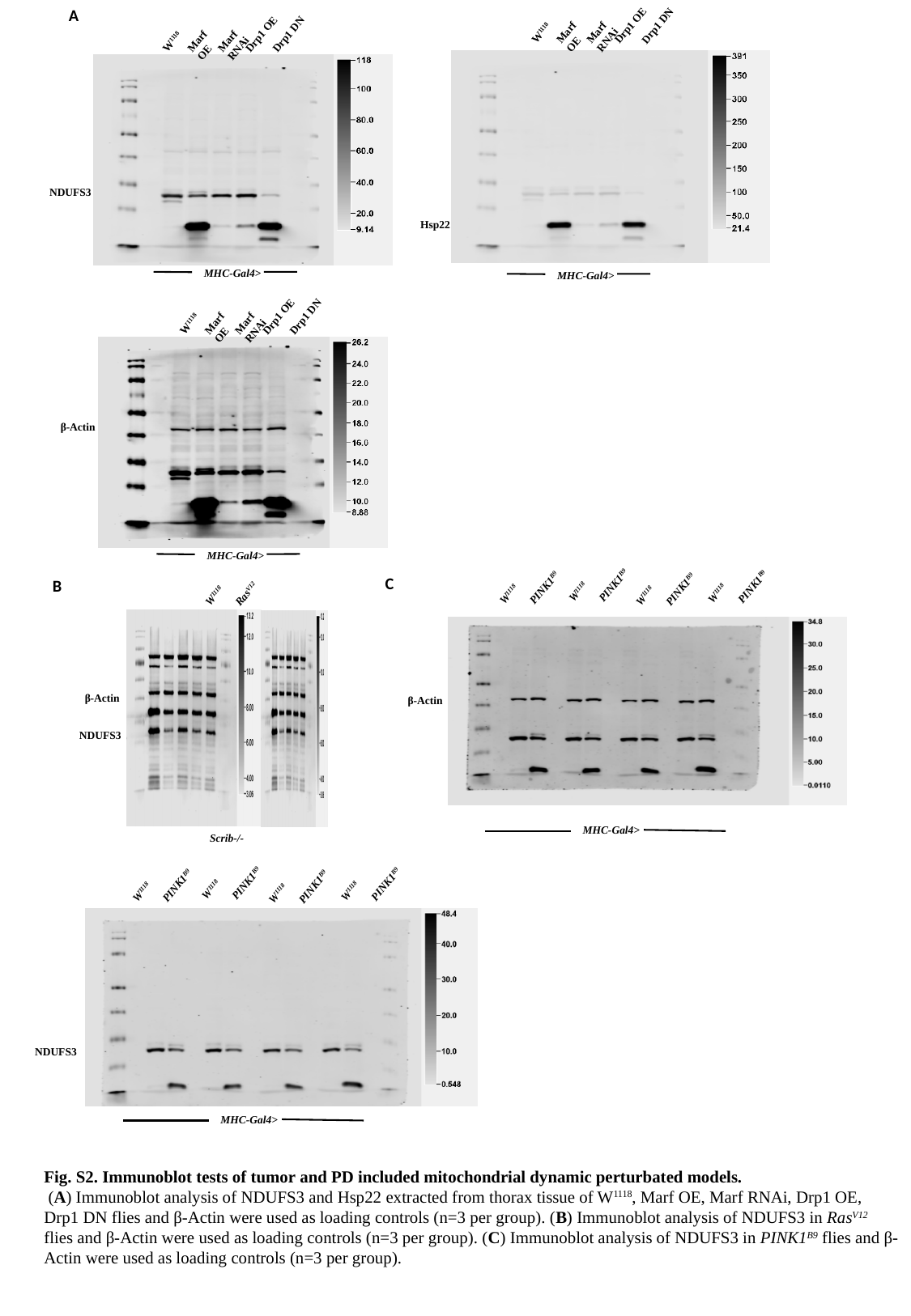

Marf RNAi
Drp1 OE
Drp1 DN
Marf OE
W^1118^
Hsp22
MHC-Gal4>
Marf RNAi
Drp1 OE
Drp1 DN
Marf OE
W^1118^
NDUFS3
MHC-Gal4>
A
Marf RNAi
Drp1 OE
Drp1 DN
Marf OE
W^1118^
β-Actin
MHC-Gal4>
PINK1^B9^
W^1118^
PINK1^B9^
W^1118^
PINK1^B9^
W^1118^
PINK1^B9^
W^1118^
β-Actin
MHC-Gal4>
Ras^V12^
W^1118^
β-Actin
NDUFS3
Scrib-/-
C
B
PINK1^B9^
W^1118^
PINK1^B9^
W^1118^
PINK1^B9^
W^1118^
PINK1^B9^
W^1118^
NDUFS3
MHC-Gal4>
Fig. S2. Immunoblot tests of tumor and PD included mitochondrial dynamic perturbated models.
 (A) Immunoblot analysis of NDUFS3 and Hsp22 extracted from thorax tissue of W^1118^, Marf OE, Marf RNAi, Drp1 OE, Drp1 DN flies and β-Actin were used as loading controls (n=3 per group). (B) Immunoblot analysis of NDUFS3 in Ras^V12^ flies and β-Actin were used as loading controls (n=3 per group). (C) Immunoblot analysis of NDUFS3 in PINK1^B9^ flies and β-Actin were used as loading controls (n=3 per group).
