## Supplementary material for "Hsp22 is the key sensor and balancer in mitochondrial dynamic associated metabolic reprogramming": Supp3

### Slide 1
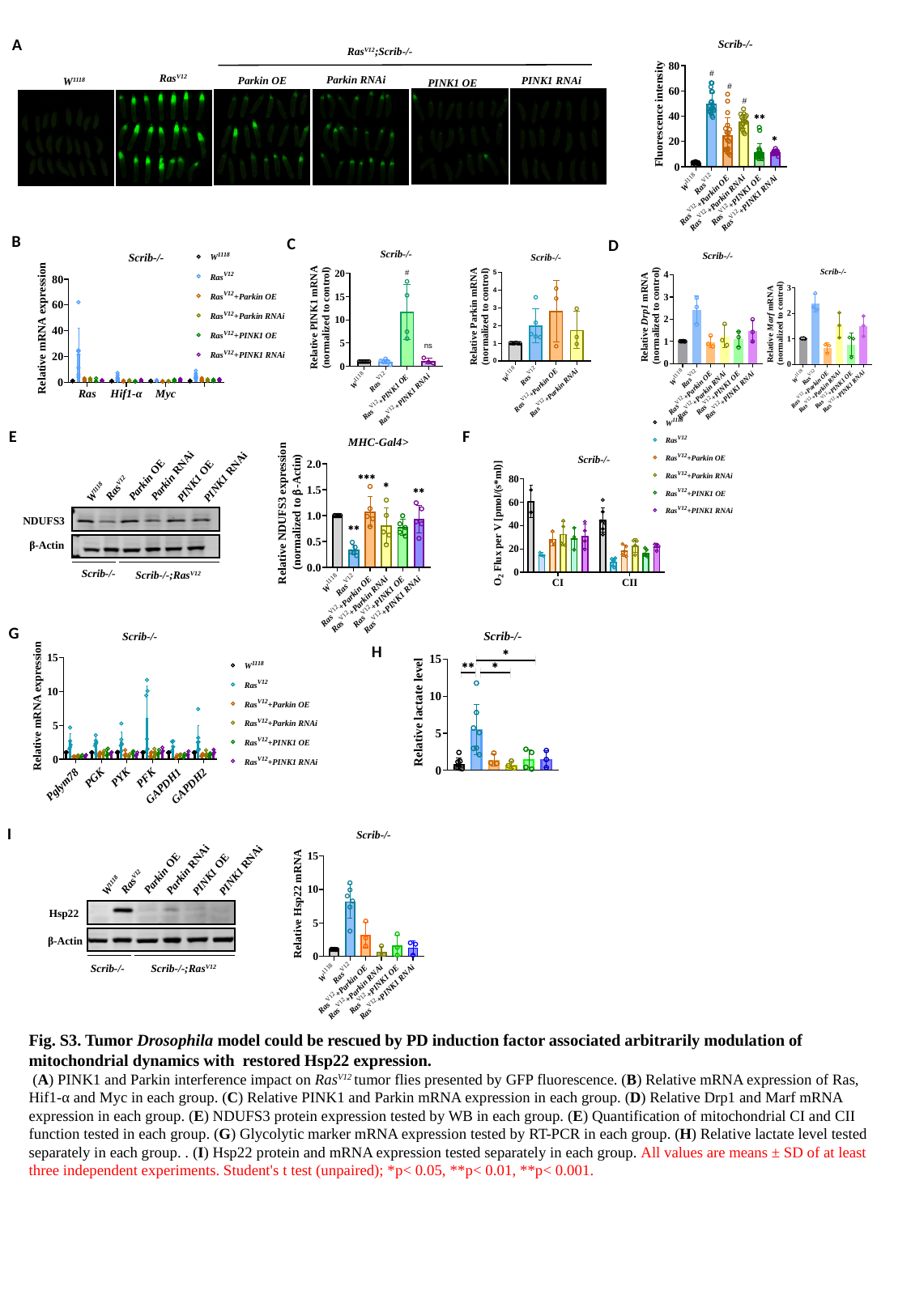

A
Ras^V12^;Scrib-/-
Ras^V12^
Parkin RNAi
Parkin OE
PINK1 RNAi
W^1118^
PINK1 OE
B
C
D
PINK1 RNAi
Parkin RNAi
Parkin OE
PINK1 OE
Ras^V12^
W^1118^
NDUFS3
β-Actin
Scrib-/-
Scrib-/-;Ras^V12^
E
F
G
H
PINK1 RNAi
Parkin RNAi
Parkin OE
PINK1 OE
Ras^V12^
W^1118^
Hsp22
β-Actin
Scrib-/-
Scrib-/-;Ras^V12^
I
Fig. S3. Tumor Drosophila model could be rescued by PD induction factor associated arbitrarily modulation of mitochondrial dynamics with restored Hsp22 expression.
 (A) PINK1 and Parkin interference impact on Ras^V12^ tumor flies presented by GFP fluorescence. (B) Relative mRNA expression of Ras, Hif1-α and Myc in each group. (C) Relative PINK1 and Parkin mRNA expression in each group. (D) Relative Drp1 and Marf mRNA expression in each group. (E) NDUFS3 protein expression tested by WB in each group. (E) Quantification of mitochondrial CI and CII function tested in each group. (G) Glycolytic marker mRNA expression tested by RT-PCR in each group. (H) Relative lactate level tested separately in each group. . (I) Hsp22 protein and mRNA expression tested separately in each group. All values are means ± SD of at least three independent experiments. Student's t test (unpaired); *p< 0.05, **p< 0.01, **p< 0.001.
