## Supplementary material for "Hsp22 is the key sensor and balancer in mitochondrial dynamic associated metabolic reprogramming": Supp4

### Slide 1
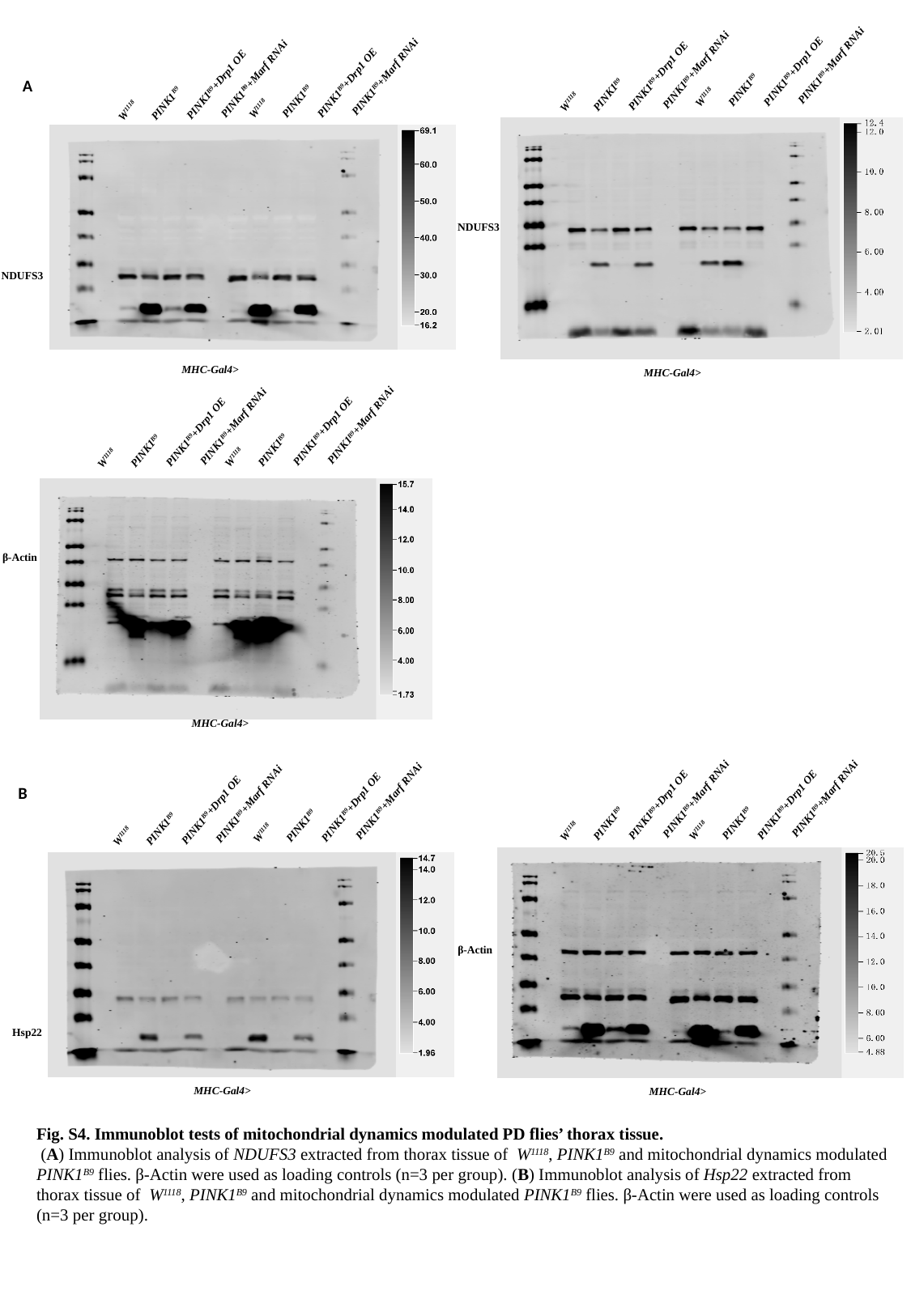

PINK1^B9^+Marf RNAi
PINK1^B9^+Drp1 OE
PINK1^B9^
W^1118^
PINK1^B9^+Marf RNAi
PINK1^B9^+Drp1 OE
PINK1^B9^
W^1118^
NDUFS3
MHC-Gal4>
PINK1^B9^+Marf RNAi
PINK1^B9^+Drp1 OE
PINK1^B9^
W^1118^
PINK1^B9^+Marf RNAi
PINK1^B9^+Drp1 OE
PINK1^B9^
W^1118^
NDUFS3
MHC-Gal4>
A
PINK1^B9^+Marf RNAi
PINK1^B9^+Drp1 OE
PINK1^B9^
W^1118^
PINK1^B9^+Marf RNAi
PINK1^B9^+Drp1 OE
PINK1^B9^
W^1118^
β-Actin
MHC-Gal4>
PINK1^B9^+Marf RNAi
PINK1^B9^+Drp1 OE
PINK1^B9^
W^1118^
PINK1^B9^+Marf RNAi
PINK1^B9^+Drp1 OE
PINK1^B9^
W^1118^
PINK1^B9^+Marf RNAi
PINK1^B9^+Drp1 OE
PINK1^B9^
W^1118^
PINK1^B9^+Marf RNAi
PINK1^B9^+Drp1 OE
PINK1^B9^
W^1118^
B
Hsp22
β-Actin
MHC-Gal4>
MHC-Gal4>
Fig. S4. Immunoblot tests of mitochondrial dynamics modulated PD flies’ thorax tissue.
 (A) Immunoblot analysis of NDUFS3 extracted from thorax tissue of W^1118^, PINK1^B9^ and mitochondrial dynamics modulated PINK1^B9^ flies. β-Actin were used as loading controls (n=3 per group). (B) Immunoblot analysis of Hsp22 extracted from thorax tissue of W^1118^, PINK1^B9^ and mitochondrial dynamics modulated PINK1^B9^ flies. β-Actin were used as loading controls (n=3 per group).
