## Supplementary material for "Hsp22 is the key sensor and balancer in mitochondrial dynamic associated metabolic reprogramming": Supp5

### Slide 1
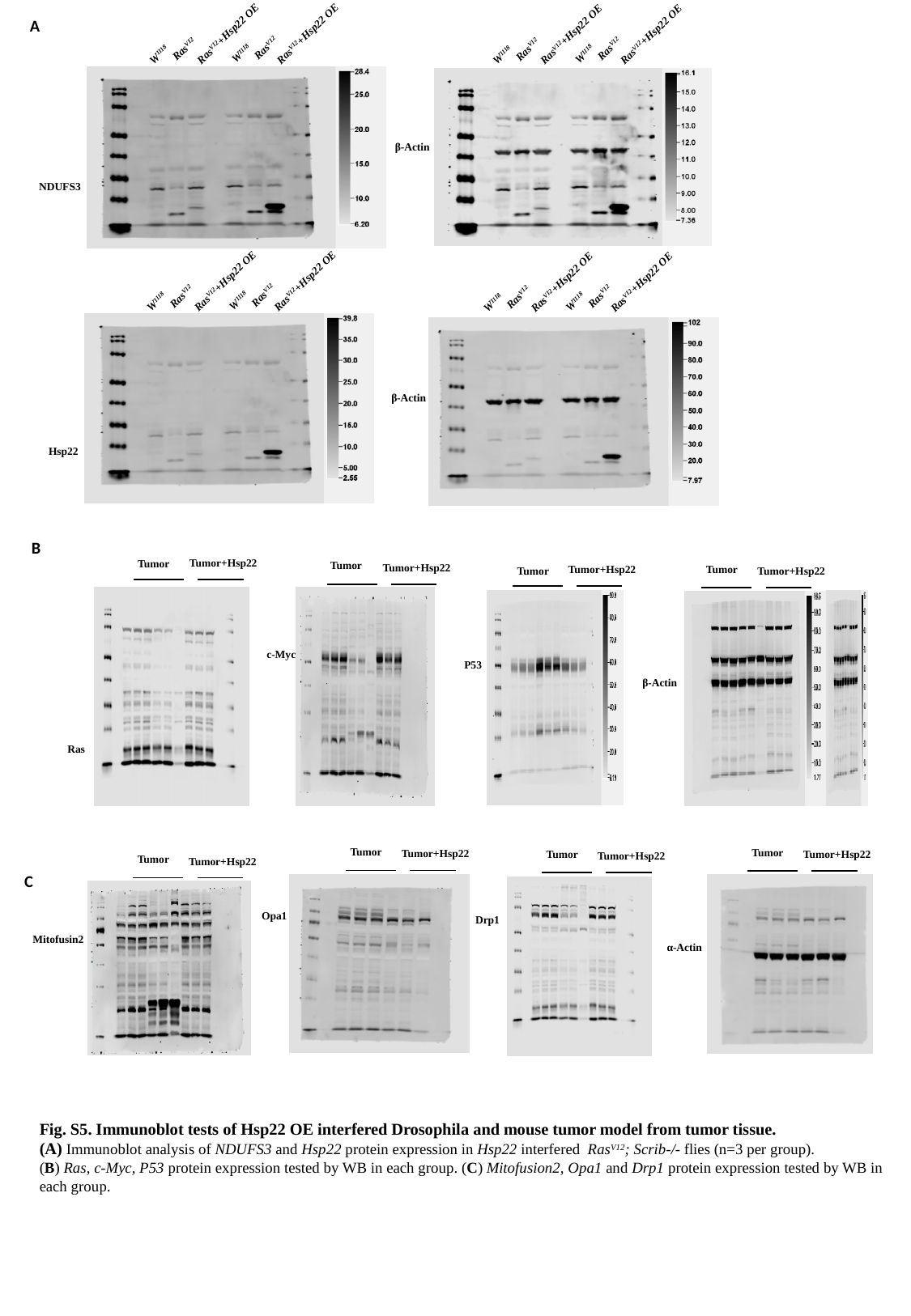

W^1118^
W^1118^
Ras^V12^+Hsp22 OE
Ras^V12^+Hsp22 OE
Ras^V12^
Ras^V12^
W^1118^
W^1118^
Ras^V12^+Hsp22 OE
Ras^V12^+Hsp22 OE
Ras^V12^
Ras^V12^
A
β-Actin
NDUFS3
W^1118^
W^1118^
Ras^V12^+Hsp22 OE
Ras^V12^+Hsp22 OE
Ras^V12^
Ras^V12^
W^1118^
W^1118^
Ras^V12^+Hsp22 OE
Ras^V12^+Hsp22 OE
Ras^V12^
Ras^V12^
Hsp22
β-Actin
B
Tumor+Hsp22
Tumor
Tumor
Tumor+Hsp22
Tumor
Tumor+Hsp22
β-Actin
Tumor+Hsp22
Tumor
Ras
c-Myc
P53
Tumor
Tumor
Tumor+Hsp22
Tumor
Tumor+Hsp22
Tumor+Hsp22
Tumor
Tumor+Hsp22
C
α-Actin
Drp1
Opa1
Mitofusin2
Fig. S5. Immunoblot tests of Hsp22 OE interfered Drosophila and mouse tumor model from tumor tissue.
(A) Immunoblot analysis of NDUFS3 and Hsp22 protein expression in Hsp22 interfered Ras^V12^; Scrib-/- flies (n=3 per group).
(B) Ras, c-Myc, P53 protein expression tested by WB in each group. (C) Mitofusion2, Opa1 and Drp1 protein expression tested by WB in each group.
