## Supplementary material for "Hsp22 is the key sensor and balancer in mitochondrial dynamic associated metabolic reprogramming": Supp6

### Slide 1
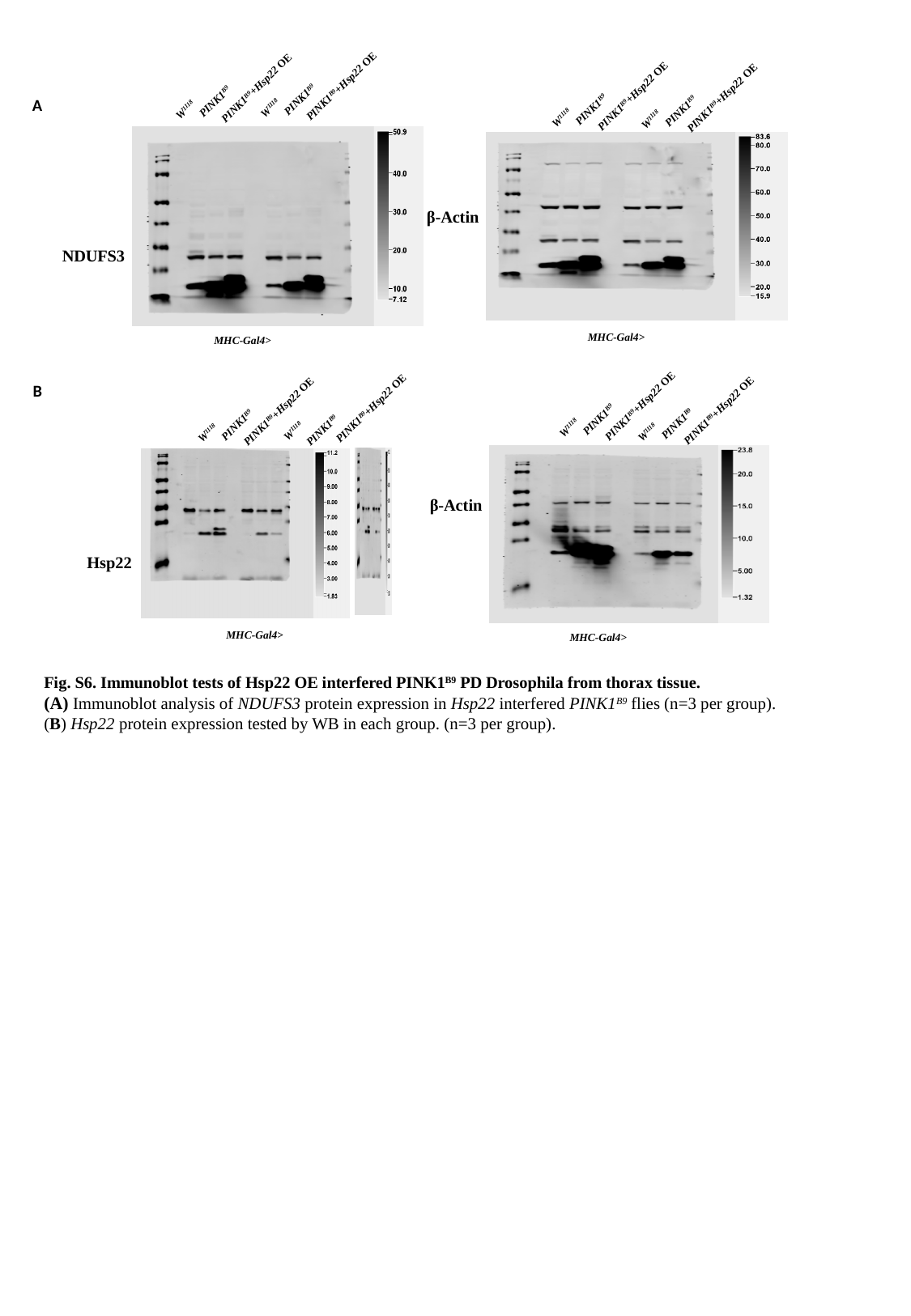

PINK1^B9^
W^1118^
PINK1^B9^+Hsp22 OE
PINK1^B9^
W^1118^
PINK1^B9^+Hsp22 OE
PINK1^B9^
W^1118^
PINK1^B9^+Hsp22 OE
PINK1^B9^
W^1118^
PINK1^B9^+Hsp22 OE
β-Actin
MHC-Gal4>
A
NDUFS3
MHC-Gal4>
PINK1^B9^
W^1118^
PINK1^B9^+Hsp22 OE
PINK1^B9^
W^1118^
PINK1^B9^+Hsp22 OE
β-Actin
MHC-Gal4>
PINK1^B9^
W^1118^
PINK1^B9^+Hsp22 OE
PINK1^B9^
W^1118^
PINK1^B9^+Hsp22 OE
Hsp22
B
MHC-Gal4>
Fig. S6. Immunoblot tests of Hsp22 OE interfered PINK1^B9^ PD Drosophila from thorax tissue.
(A) Immunoblot analysis of NDUFS3 protein expression in Hsp22 interfered PINK1^B9^ flies (n=3 per group).
(B) Hsp22 protein expression tested by WB in each group. (n=3 per group).
